# Changes to zonular tension alters the subcellular distribution of AQP5 in regions of influx and efflux of water in the rat lens

**DOI:** 10.1101/2020.06.29.178756

**Authors:** Rosica S Petrova, Nandini Bavana, Rusin Zhao, Kevin L Schey, Paul J Donaldson

**Affiliations:** Department of Physiology, School of Medical Sciences, New Zealand National Eye Centre, University of Auckland, Auckland, New Zealand; Mass Spectrometry Research Center, Vanderbilt University, Nashville, TN, USA

**Keywords:** Lens, water transport, immunohistochemistry, AQP0, AQP1, AQP5, zonular tension

## Abstract

**Purpose:** The lens utilizes circulating fluxes of ions and water that enter the lens at both poles and exit at the equator to maintain its optical properties. We have mapped the subcellular distribution of the lens aquaporins (AQP0, 1, & 5) in these water influx and efflux zones and investigated how their membrane location is affected by changes in tension applied to the lens by the zonules.

**Methods:** Immunohistochemistry using AQP antibodies was performed on axial sections obtained from rat lenses that had been removed from the eye and then fixed, or were fixed *in situ* to maintain zonular tension. Zonular tension was pharmacologically modulated by applying either tropicamide (increased), or pilocarpine (decreased). AQP labelling was visualized using confocal microscopy.

**Results:** Modulation of zonular tension had no effect on AQP1 or AQP0 labelling in either the water efflux, or influx zones. In contrast, AQP5 labelling changed from membranous to cytoplasmic in response to both mechanical and pharmacologically induced reductions in zonular tension in both the efflux zone, and anterior (but not posterior) influx zone associated with the lens sutures.

**Conclusions:** Altering zonular tension dynamically regulates the membrane trafficking of AQP5 in the efflux and anterior influx zones to potentially change the magnitude of circulating water fluxes in the lens.

## INTRODUCTION

The transparency and refractive properties of the lens are established and maintained by its unique cellular structure and function^1^. Structurally, the lens is attached to the ciliary body via the lens zonules, and is bathed on its anterior and posterior surfaces by the aqueous and vitreous humors, from which it exchanges nutrients and waste products. Underneath the capsule the anterior surface of the lens is covered by a single layer of epithelial cells that, at the equatorial margins, divide and differentiate in the lens fiber cells which make up the bulk of the lens (Figure 1A). These fiber cells undergo a process of differentiation that involves not only a change of the molecular profile of the newly differentiated fiber cells, but also a massive elongation of their lateral membrane domains towards the poles of the lens. This process continues until either the apical or basal membrane tips of the fibers meet with the membrane tips of fibers cells from adjacent hemisphere to form the anterior and posterior sutures, respectively^2^. As fiber cells differentiate, they lose their light scattering organelles and internalize older fiber cells into the lens core or nucleus. Since this process occurs throughout life a gradient of fiber cell age is established with oldest mature fiber cells in the lens nucleus having been laid down during embryogenesis. Since the lens lacks a blood supply, it instead operates an internal microcirculation system (Figure 1A), that acts to maintain the overall tissue architecture of the lens by delivering nutrients, removing waste products, and controlling the volume of lens fiber cells^3^. This internal microcirculation is generated by circulating ionic and fluid fluxes that enter the lens at both poles via an extracellular pathway, associated with the lens sutures, which directs ions and water towards the deeper regions of the lens (Figure 1A). Ions and water then cross fiber cell membranes before returning to the lens surface via an intercellular outflow pathway mediated by gap junctions. The gap junctions direct the outflow of ions and water to the lens equator where the Na^+^ pumps, which generate the circulating flux of Na^+^ ions that drive the microcirculation system, and other ion and water channels are localized allowing ions and water to cross the membranes of surface cells and leave the lens.

**Figure 1.**
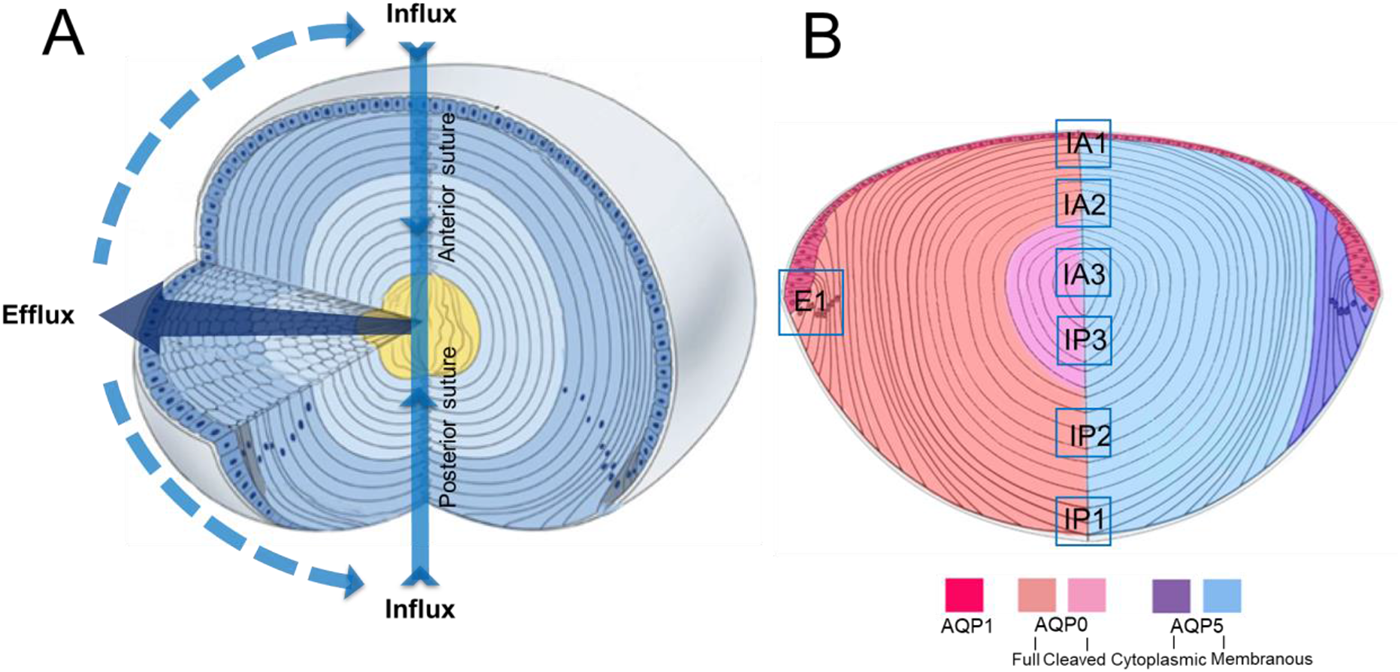
Schematic diagrams illustrating lens structure and function and the distribution of AQPs in different regions of the lens. (A) 3D diagram of the lens showing ion and water fluxes coming into the lens core (*yellow*) via an extracellular route located at the anterior and posterior sutures (*blue arrows*). Ions and water cross fiber cell membranes before travelling via an intercellular pathway mediated by gap junction channels (*dark blue arrow*) to exit the lens at the equator. (B) Diagram of an axial section of the lens showing subcellular distributions of the lens water channels (AQP) in the different regions of the lens. AQP1 (*red*) is restricted to the membranes of the lens epithelium. AQP0 (*left*) is found in the membranes of lens fiber cells across all areas of the lens, but in the core of the lens the C-terminal tail is cleaved. AQP5 (*right*) is also found throughout all regions of the lens, but in the epithelial (*not shown*) and peripheral differentiating fiber (*purple*) cells it is associated with the cytoplasm. In deeper regions of the outer cortex AQP5 becomes associated with the plasma membrane (*blue*), and this labelling extends into the lens core. In this study we focused on the subcellular distributions of the three lens AQPs at the equator, which is associated with water efflux (E1), and the anterior (IA1, IA2, IA3) and posterior (IP1, IP2, IP3) poles which mediate water influx. Adapted with permission from Shi *et al*. (2009).

The intercellular outflow of water through the gap junction generates a substantial hydrostatic pressure gradient^4^, that is not only remarkably similar amongst different species of lens^5^, but it is also highly regulated by a dual feedback pathway^6^. This pathway utilizes the mechanosensitive non-selective ion channels TRPV1 and TRPV4 to sense changes in lens pressure and to activate signaling pathways that reciprocally modulate the activity of Na^+^ pumps at the lens equator to ensure the pressure gradient is maintained constant. More recently, Chen et al., have shown that altering the zonular tension applied to the lens can alter this constant hydrostatic pressure set point^7^. Thus, while it is apparent that alterations in ionic/osmotic gradients, via modulation of Na^+^ pump activity, changes the flow of water (pressure) through the lens, whether the permeability of lens cells to water is also modulated in parallel to changes in the osmotic gradient has yet to be determined.

As in other tissues, the water permeability (P_H2O_) of lens cell membranes is determined by the profile of the different aquaporin (AQP) proteins that are expressed in the different regions of the lens^8^. Three different water channels, AQP0, AQP1 and AQP5 (Figure 1B), all with very different P_H2O_ and regulatory properties are differentially expressed in the lens^9–11^. AQP1, a constitutively active water channel with a high P_H2O_, is found only in the epithelium^11, 12^. AQP0, the most abundant membrane protein in the lens, is found only in the fiber cells^13^. AQP0 is actually a relative poor water channel^9, 14, 15^, but has been shown to have additional roles as an adhesion protein^16–18^ and a scaffolding protein^19, 20^, and undergoes extensive post-translational cleavage of its C-terminal tail in mature fiber cells^15, 21, 22^. AQP5, in other tissues, has been shown to act as a regulated water channel with a relatively high P_H2O_, which when inserted into the membrane increases the P_H2O_ of epithelial cells^23–26^. In the lens, immunohistochemical mapping of the subcellular distribution of AQP5 has shown it to be expressed in both the epithelium and throughout the fiber cell mass^22, 27^. In peripheral fiber cells it is initially found as a cytoplasmic pool of protein that undergoes a transition to the membrane as fiber cells differentiate and become internalized. However, unlike AQP0, AQP5 does not undergo extensive cleavage of its C-terminus in the lens nucleus^22^.

In this study, we have mapped the subcellular distribution of the three lens AQPs to determine whether changes in P_H2O_ contribute to the observed modulation of water transport. We have focused on their relative distributions at the equator and both poles, the regions of the lens involved in water efflux and influx, respectively. Furthermore, in an effort to induce changes in P_H2O_ we have modified zonular tension, which alters hydrostatic pressure^7^, and have used immunohistochemistry to visualize induced changes in the subcellular location of the AQPs in the water efflux and influx zones. Our results suggest that dynamic changes in the subcellular location of AQP5 may differentially alter the P_H2O_ of fiber cell membranes in the efflux and influx zone and therefore may contribute to the regulation of water transport in the lens.

## METHODS

### Reagents

A polyclonal affinity purified anti-AQP0 antibody, directed against the last 17 amino acids of the C-terminus of the human protein, was obtained from Alpha Diagnostic International, Inc. (San Antonio, TX, USA, catalogue number AQP01-A). An affinity purified polyclonal anti-AQP5 antibody, directed against the last 17 amino acids of the C-terminus of the rat protein, was obtained from Merck Millipore (Darmstadt, Germany, catalogue number AB15858). A rabbit polyclonal antibody directed against the last 19 amino acids of the C-terminus of the rat AQP1 protein was purchased from Alpha Diagnostic International (San Antonio, Texas, USA). Secondary antibodies (goat anti-Rabbit Alexa 488), and wheat germ agglutinin (WGA) conjugated to a fluorophore (WGA-Alexa 594) for labelling of the cell membrane, were obtained from Thermo Fisher Scientific (Waltham, MA). For labelling of the epithelial and fiber cell nuclei, DAPI was obtained from Sigma–Aldrich (St Louis, MO, USA). Phosphate buffered saline (PBS) was prepared fresh from PBS tablets. Lenses were organ cultured in Artificial Aqueous Humour (AAH) that consisted of (in mM): 125 NaCl, 0.5 MgCl_2_, 4.5 KCL, 10 NaHCO_3_, 2 CaCl_2_, 5 Glucose, 10 sucrose, 10 HEPES, pH 7.4, 300mOsml/L. Tropicamide and pilocarpine were used at 1:5 dilution prepared in AAH buffer from 1% w/v eye drops. Unless otherwise stated all other chemicals were from Sigma-Aldrich

### Lens organ cultures

All animal experiments were carried out in accordance with the ARVO Statement for the Use of Animals in Ophthalmic and Vision Research and were approved by the University of Auckland Animal Ethics Committee (# 001893). Eyes were enucleated from 21-day-old Wistar rats and subjected to three different preparation protocols that yielded lenses that were either separated from, or attached to, the ciliary muscle by the zonules. In the first preparation, lenses were separated from the ciliary body by completely removing the lens from the eye. To achieve this, four radiating incisions from the optic nerve head towards the limbus were first made to expose the lens and ciliary body. To release the lens, the zonules were cut with a pair of surgical scissors, the lens was removed from the eye using a glass loop, and the excised lens was organ cultured in AAH for up to 120 minutes before immersing in 0.75% PFA prepared in PBS pH 7.4 and fixed overnight at room temperature. In the second preparation, lenses were prepared with the zonules attached by first cutting a small (~2 mm) hole into the cornea of the enucleated eye, to provide a pathway to perfuse the lens with AAH in the absence or presence of either tropicamide or pilocarpine. Lenses were maintained in organ culture for up 120 minutes at 37°C in a CO_2_ incubator (Heracell 150i, Thermo Scientific, USA). Tropicamide causes a relaxation of the ciliary muscle and increases the tension applied to the lens via the zonules, while pilocarpine by inducing a contraction of the ciliary muscle reduces the tension applied to the lens via the zonules. After organ culture, the preparation was perfused with 0.75% PFA, and maintained at room temperature overnight to fix the lens *in situ*. After fixation lens were dissected free from the surrounding tissue and processed for immunohistochemistry.

The third preparation was used to confirm the ability of tropicamide and pilocarpine to alter zonular tension. The drugs were diluted in AAH and applied to the corneal surface of enucleated eyes, and the native three-dimensional structure of the ciliary body, zonules and lens, was preserved by fixing the eyes in 4% paraformaldehyde, while maintaining the intraocular pressure at 18 mmHg^28^ using the method described by Bassnett^29^. After fixation, the posterior sclera and retina were removed, and the ciliary body and lens photographed using a stereo microscope (Leica Microsystems, Germany). The distance between the two structures was measured using image analysis software (Adobe Photoshop CC). For each lens ~25 measurements were taken to calculate the mean circumferential space distance from at least three different lenses incubated in the absence and presence of either tropicamide or pilocarpine.

### Immunohistochemistry

Fixed lenses were washed 3 times in PBS pH 7.4 for 10 minutes, and cryoprotected using 3 consecutive incubations in 10% and 20% sucrose for 1 hour at room temperature, followed by an overnight incubation in 30% sucrose at 4°C^30^. Cryoprotected lenses were positioned on chucks in an axial orientation, encased in optimal temperature medium (OCT, Japan), and snap frozen for 15 seconds in liquid nitrogen. Axial sections were cut using a Leica CM3050S cryostat (Leica Biosystems, Germany), and consecutive sections of ~14 μm thickness were collected from at least three lenses for each experimental condition to ensure consistency of immunolabelling results. Sections were washed 3 times for 5 minutes in PBS pH 7.4 and blocked for 1 hour in blocking solution (3% normal goat serum, 3% BSA dissolved in PBS, pH 7.4). Excess blocking solution first was removed using tissue paper, before incubating sections in different anti-AQP primary antibodies diluted 1:100 in blocking solution overnight at 4°C. Sections were washed 3 times for 5 minutes in PBS pH 7.4, to remove unbound antibody, before incubating sections with a goat anti-rabbit Alexa 488 secondary antibody for 3 hours at room temperature. After washing the secondary antibody 3 times in PBS pH 7.4 for 5 minutes, sections were incubated at room temperature for 1 hour in the membrane marker WGA and the nucleus marker DAPI (0.25μg/ml), diluted 1:100 and 1:1000 in PBS pH 7.4, respectively. After a final wash in PBS pH 7.4, sections were mounted with a cover slip using an anti-fading agent (Vectashield, Vector Laboratories, San Diego, CA, USA), and imaged using an Olympus FV1000 Fluoview confocal scanning microscope (Tokyo, Japan). The resultant images were prepared using Adobe Photoshop CC software.

### Statistical Analysis

Experimental means are given as ± standard error of the mean (SE). Statistical significance was tested with the Mann-Whitney *U*-test, using GraphPad Prism (La Jolla, CA). Statistical significance was set at the α = 0.05 level.

## RESULTS

In previous studies we have mapped the distribution of AQP0^21^ and AQP5^22^ from the periphery to the center of the lens using sections taken through the equator of the rodent lenses, which were removed from the eye by cutting the zonules that attach the lens to the ciliary body. In this study, we have utilized axial sections to map the subcellular distribution of AQPs in the rat lens, as this orientation allows us to visualize, in the same section, the anterior and posterior poles of the lens that mediate water influx, as well as the lens equator where water efflux occurs (Figure 1). Furthermore, to determine the effects of changes in zonular tension on the subcellular distribution of AQPs, axial sections were obtained from lenses that had been subjected to mechanical and pharmacological manipulations to alter the tension applied to the lens via the zonules. The effects of altering zonular tension on the subcellular localization of the three major AQPs 1, 0, and 5 in the water efflux and influx pathways are presented in turn.

### The subcellular distribution of lens AQPs in the equatorial water efflux zone

As epithelial cells at the lens equator initiate the process of differentiation into fiber cells in the water efflux zone, they change their AQP expression profile (Figure 2). As had been shown previously in lenses removed from the eye by cutting the zonules, AQP1 was expressed only in the epithelial cells that cover the anterior surface of the lens^12^, with AQP1 labelling being strongly localized to the apical and lateral membrane domains of the cells (Figure 2B). However, as the epithelial cells elongated into fiber cells, AQP1 labelling abruptly disappeared and was completely lacking in secondary fiber cells (Figure 2B). In contrast to AQP1, AQP0 protein was not detected in epithelial cells, and was initially only observed in the newly-derived elongating fiber cells as a cytoplasmic punctate labelling pattern, with strong membranous labelling only becoming apparent in fiber cells ~20-25 cell layers in from the capsule (Figure 2C). AQP5 labelling was initially predominately cytoplasmic in both the epithelial and newly differentiated fiber cells (Figure 2D), before becoming localized to the membranes of secondary fiber cells ~150 to 200μm in from the capsule in an area just past the bow region where cell nuclei disperse (Figure 2E). In summary, we see a change in expression from AQP1 to AQP0 as epithelial cells differentiate into fiber cells, while AQP5 is expressed in both cell types. In the efflux zone, both AQP1 and AQP0 show an exclusive membrane localization that suggests they both contribute to P_H2O_ in this zone, while, the localization of AQP5 will the cytoplasm indicates AQP5 is not actively contributing to the P_H2O_ of epithelial cells and peripheral fiber cells in the efflux zone. However, in deeper differentiating fiber cells the insert of AQP5 in the plasma membrane indicates that in these cells it does contribute to P_H2O_^27^.

**Figure 2.**
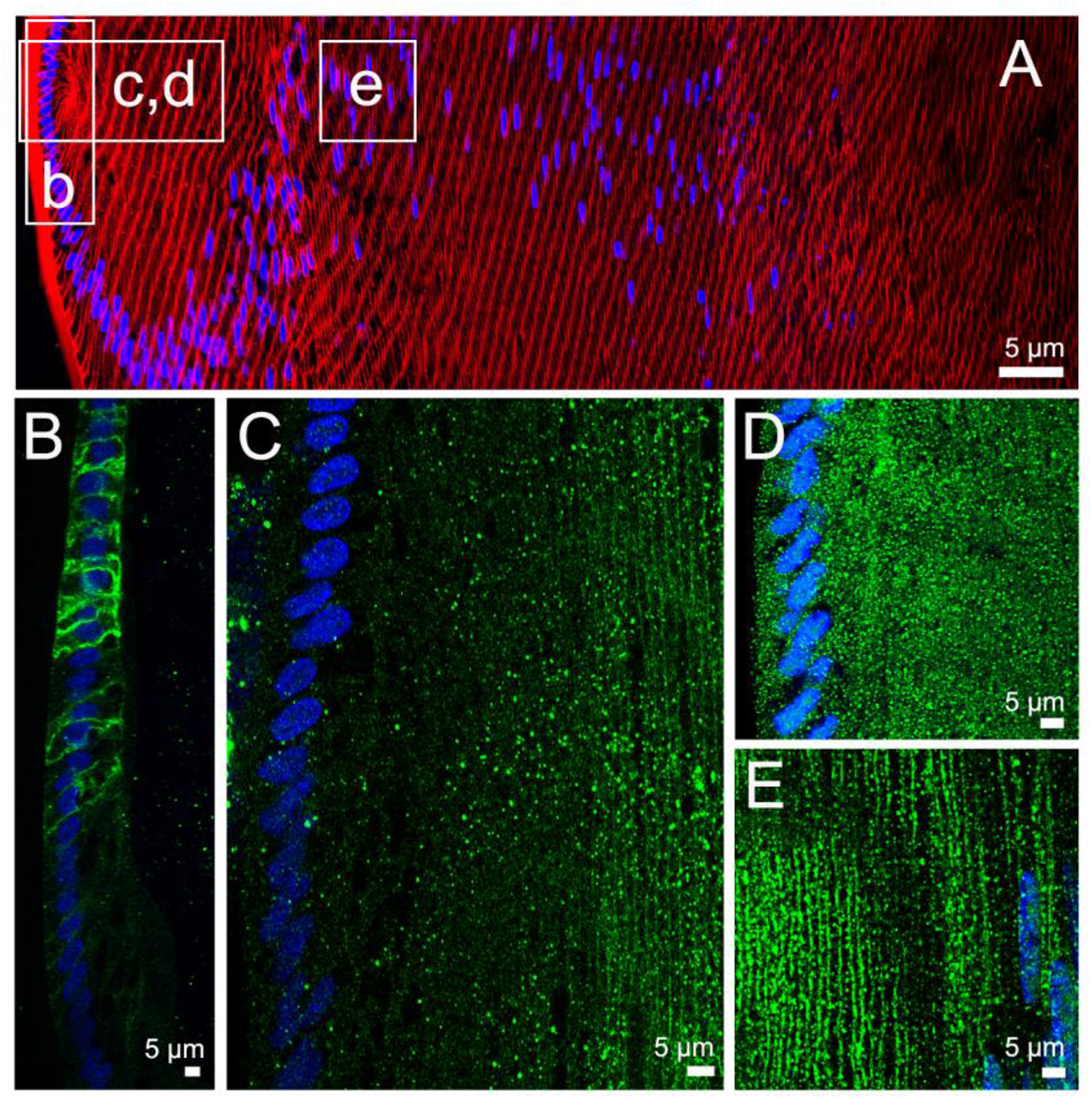
Subcellular localization of lens AQPs in the efflux zone of rat lenses in the absence of zonular tension. (A) Image montage of the water efflux zone taken from an axial section of a rat lenses that had been removed from the eye by first cutting the zonules, and was then placed immediately into fixative. The section was labelled with the membrane marker WGA (*red*) and the nuclei marker DAPI (*blue*). Boxes indicate that areas from which the higher resolution images shown in images B-E were taken from to investigate the subcellular distribution for each lens AQP (*green*). (B) AQP1 labelling is localized to the membranes of the epithelial cells but disappears as the epithelial cells differentiate into fiber cells. (C) AQP0 labelling was absent from epithelial cells, but became apparently initially as diffuse punctate labelling as epithelial cell differentiated into fiber cells, and became strongly associated with the membranes of fiber cells are later stages of differentiation deeper localized secondary fiber cells. (D) AQP5 labeling was initially strongly cytoplasmic in the epithelial cells and newly differentiated fiber cells and only became membranous in differentiating fiber cells at ~150 μm distance from the capsule of the lens (E).

To determine whether altering zonular tension changes the subcellular distribution of the lens AQPs, we organ cultured rat lenses either *ex vivo* (no zonules) or *in situ* (with zonules) for different periods of time before fixing the lenses for immunohistochemistry. The pattern of AQP1 (Figure 2B) and AQP0 (Figure 2C) labelling observed in lenses organ cultured in lenses with no zonules attached was not altered in lenses with zonules attached (data not shown). However, the subcellular labelling patterns observed for AQP5 were different in lenses maintained in *ex vivo* or *in situ* organ culture (Figure 3). Lenses, maintained in *ex vivo* organ culture after having their zonules cut, showed an initial predominantly cytoplasmic subcellular localization of AQP5 in both epithelial and fiber cells for up to the first 45 minutes in culture. Then, over the next 60 to 120 minutes the labelling became increasingly associated with the plasma membrane of peripheral fiber cells in the water efflux zone (Figure 3B). In contrast, lenses that were fixed *in situ*, with their zonules intact, exhibited predominately membranous AQP5 labelling in fiber cells throughout the efflux zone that did not change during the entire 120 minutes of organ culture (Figure 3C). Taken together these results suggest that AQP5 in the water efflux zone is normally membranous and that reducing zonular tension results in the rapid removal of AQP5 from the membrane.

**Figure 3.**
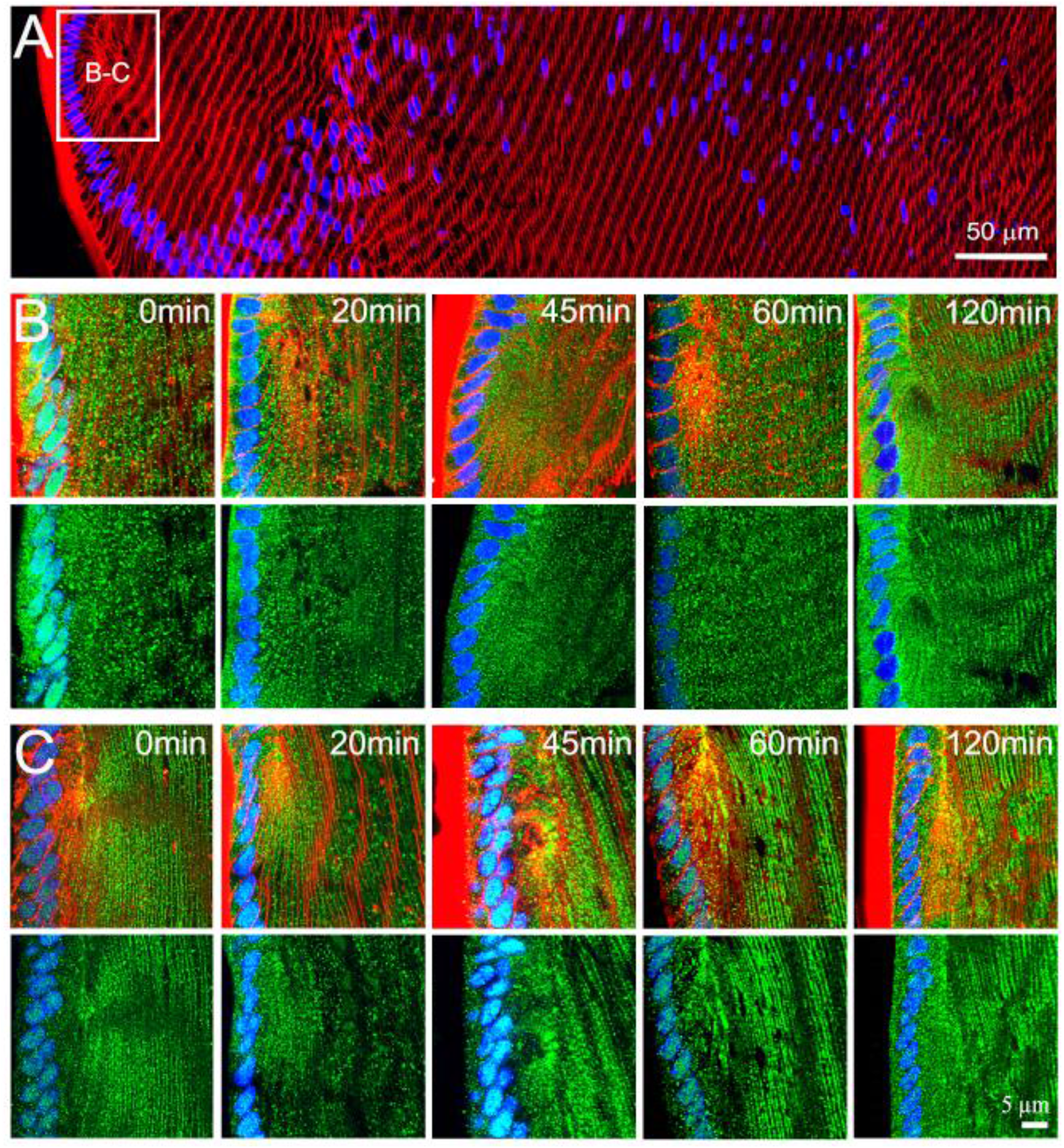
Effects of mechanically altering zonular tension on the subcellular localization of AQP5 in the efflux zone of the rat lens. (A) Image montage of the water efflux zone taken from a representative axial section of a rat lenses labelled with the membrane marker WGA (*red*) and the nuclei marker DAPI (*blue*). The box indicates the area from which high resolution images (B & C) were captured to monitor the time course of changes to the subcellular distribution of AQP5 (*green*) over a period of up to 120 minutes, in lenses that had been removed from the eye by cutting the zonules (B), or in lenses maintained *in situ* with their zonules intact (C). *Top panels* in B & C show nuclei, membrane and AQP5 labelling, while bottom panels show only nuclei and AQP5 labelling. In lenses maintained in organ culture with their zonules cut (B) the subcellular localization of AQP5 changes from a cytoplasmic to a membranous labelling pattern over time. While in lenses in which the zonular tension is maintained (C), AQP5 labelling is associated with the membrane and this labelling does not change over time in organ culture.

To further test this notion, we incubated enucleated eyes in the absence or presence of either tropicamide or pilocarpine, to pharmacologically manipulate iris and ciliary muscle contractility, in order to alter the tension applied to the lens *in situ* via the zonules (Figure 4). As expected, tropicamide (Figure 4B) and pilocarpine (Figure 4C) evoked dilation and constriction, respectively, of the pupil of the enucleated rat eye, confirming the functionality of drugs on the muscles of the iris. To assess the functionality of the drugs on the ciliary muscle, we first fixed the entire globe, before removing the posterior tissues of the eye to reveal the ciliary body and lens in order to allow the distance between the ciliary process and lens, the circumlental space, to be measured. Under control conditions the circumlental space was 151.51 + 2.3 μm (n = 5), which was significantly increased in the presence of tropicamide by 11% to 167.67 + 1.4 μm (n = 3), and significantly decreased by 14% to 129.03 + 2.5 μm after incubation in pilocarpine (n = 3) (Figure 4H). Having demonstrated that we could pharmacologically modulate ciliary muscle contractility to alter the distance between the ciliary processes and the lens, we used these pharmacological tools to alter the tension applied to the lens via the zonules.

**Figure 4.**
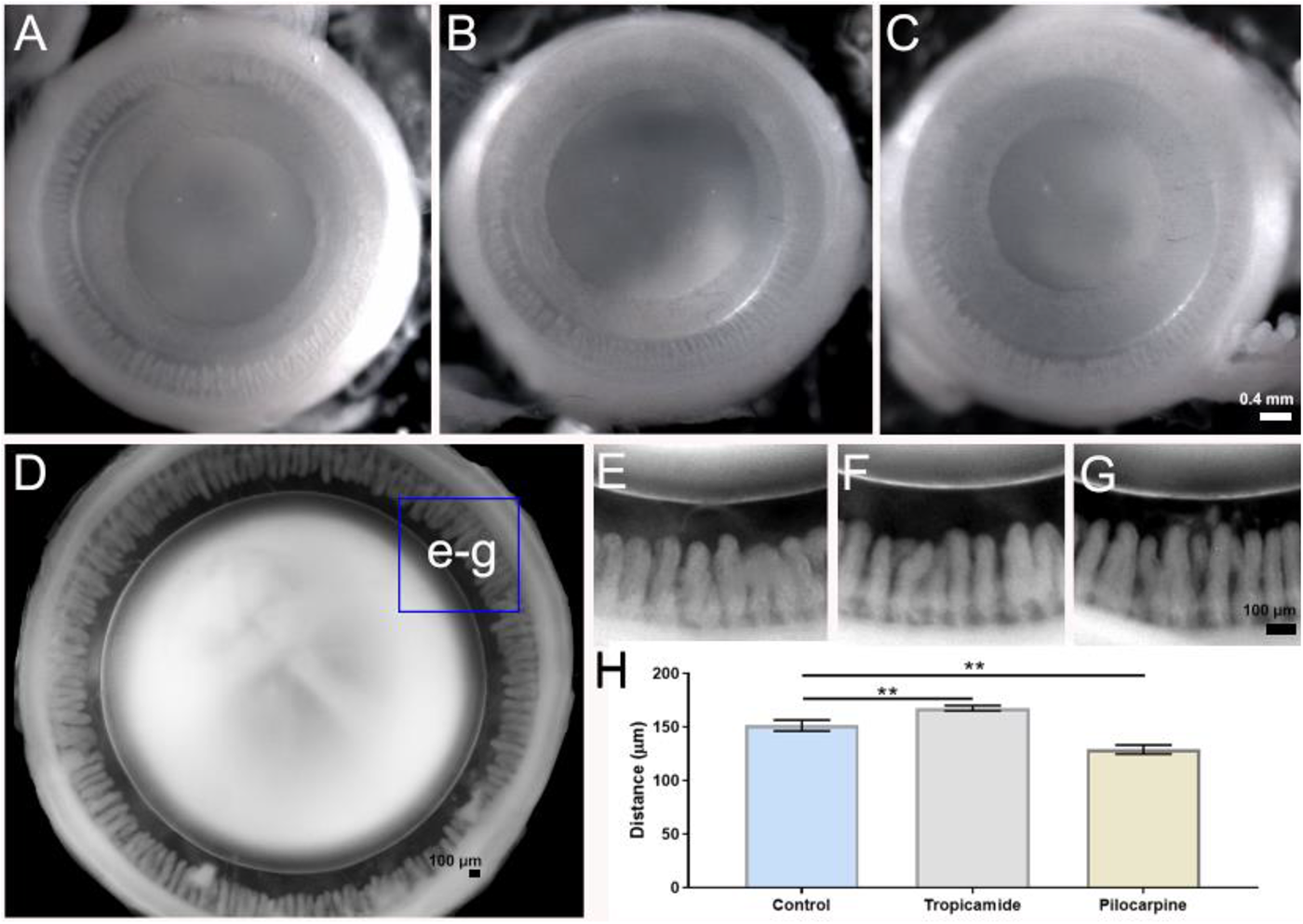
Pharmacological modulation of zonular tension of the rat lens. (A-C) Images looking down on the anterior surface of enucleated rat eyes showing the pupil diameter in untreated eyes (A), and the increase and decrease following treatment with either tropicamide (B) or pilocarpine (C), respectively, for 60 minutes. (D-G) Eyes were then fixed and the posterior sclera and retina removed to visualize the circumlental space (D&E) and how it increased and decreased following treatment with either tropicamide (F) or pilocarpine (G). (H) Summary of measurements taken from high power images showed that in control eyes (E), the distance between ciliary processes and the lens was 151.51 ± 2.3 μm (mean ± SE). In eyes treated with 0.2% tropicamide (J), the circumlental space was increased to 167.67 ± 1.4 μm (mean ± SE). In eyes treated with 0.2% pilocarpine (I), the circumlental space was reduced to 129.03 ± 2.5 μm (mean ± SE). These differences were statistically significant (P < 0.05). Statistical analysis was performed with a Mann-Whitney U test, P < 0.05 for each tested group.

If the assumption that ciliary muscle contractility alters zonular tension in our experimental model is true, we would expect that narrowing of the circumlental space induced by pilocarpine should have a similar effect on the subcellular distribution of AQP5 as that seen in lenses fixed immediately after having their zonules mechanically cut to release the tension applied to the lens. To test this notion immunohistochemistry for AQP5 was performed on lenses fixed *in situ* following incubation in the absence or presence of either tropicamide, or pilocarpine (Figure 5). We found that increasing the tension applied to the lens via the zonules by the application of tropicamide had no effect on the membrane location of AQP5 (Figure 5C), compared to control lenses with intact zonules (Figure 5B). In contrast, eyes incubated in pilocarpine that reduces the tension applied to the lens exhibited AQP5 labelling that was predominately associated with the cytoplasm of fiber cells in the water efflux zone (Figure 5D). In summary, it appears that both mechanically and pharmacologically reducing the tension applied to the lens switches AQP5 labelling from the membrane to cytoplasm, which suggests AQP5 functionality can be dynamically regulated to alter water efflux at the lens equator. In the next sections we investigate whether similar changes in the subcellular distribution of lens AQPs occurred in the water influx zones located at the anterior and posterior poles.

**Figure 5.**
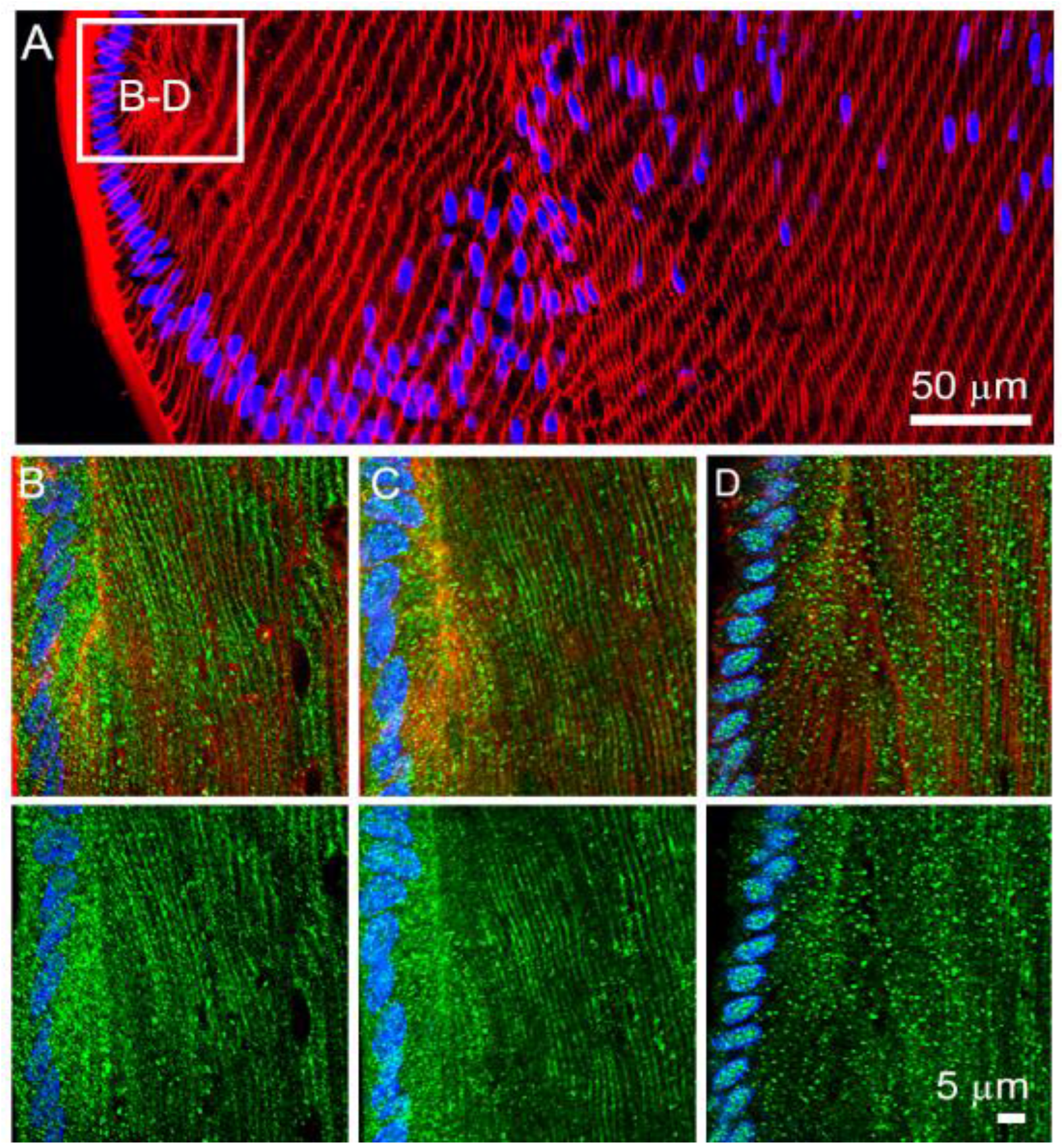
Effects of pharmacologically altering zonular tension on the subcellular localization of AQP5 in the efflux zone of the rat lens. (A) Image montage of the water efflux zone taken from a representative axial section of a rat lenses labelled with the membrane marker WGA (*red*) and the nuclei marker DAPI (*blue*). The box indicates the area from which high resolution images (B-D) were captured in lenses maintained *in situ* with their zonules intact. (B-D) *Top panels* show nuclei, membrane and AQP5 labelling, while bottom panels show only nuclei and AQP5 labelling from control lenses (B), and lenses incubated in tropicamide (C), or pilocarpine (D) for 60 minutes. Note the shift from membranous labelling to cytoplasmic labelling following incubation on pilocarpine.

### The subcellular distribution of lens AQPs in the water influx zone

Experiments that have utilized Ussing chambers^31^ and MRI measurements of heavy water penetration into the lens^32^, have shown that water preferentially enters the lens at its anterior and posterior poles. At the poles, fiber cells from adjacent hemispheres meet to form the lens sutures^2^, which in the rat lens form a Y-shaped structure. The sutures form an extracellular pathway that links the mature fiber cells in the center of the lens to the aqueous and vitreous humors that bathe the anterior and posterior poles of the lens, respectively. As such the sutures represent a pathway to direct ion and fluid movement towards the internalized mature fiber cells^33^. In axial sections it is possible to visualize the sutures as a line extending from the surface to the core of the lens, but it is difficult to observe the anterior and posterior sutures in the same section as they are offset from each other by 60°^34^. In this study, we have investigated the distribution of the AQPs in anterior (IA) and posterior (IP) influx zones by mapping the subcellular distribution of the three AQPs along the two suture lines that extend from the anterior and posterior poles to the lens core. To facilitate this comparison we have designated three spatial regions in the outer cortex (IA1; IP1), inner cortex (IA2; IP2) and core (IA3; IP3) from which higher resolution images have been taken to enable comparison between the labelling obtained from lenses exposed to different degrees of zonular tension (Figure 1B). Results from the anterior and posterior poles are presented in turn.

#### AQP distributions along the posterior suture

Since AQP1 is not present in the fiber cells below the germinative zone, we only examined the distribution of AQP0 and AQP5 in the posterior influx zone. We found that in lenses with their zonules cut, AQP0 was localized to the lateral membranes of fiber cells and was concentrated in the basal tips of fibers that interact to form the sutures in the outer cortex of the lens (Figure 6B, IP1). In contrast, AQP5 was cytoplasmic with intermittently sparse clusters of puncta on the lateral membranes of fiber cells, but no localization of AQP5 was observed at the basal tips of fiber cells that interface to form the posterior suture (Figure 6C, IP1). In the inner cortex AQP0 remained membranous and associated with the suture, while AQP5 was predominately associated with lateral membranes, but not the posterior suture (Figure 6B&C, IP2). In mature fiber cells located in the lens core AQP0 labelling was abolished, as expected by the C-terminal cleavage of the AQP0 protein as AQP0 protein undergoes a cleavage of its cytoplasmic C-terminus tail, which removes the epitope detected by the AQP0 antibody used in this study^21^. In contrast, strong AQP5 labelling was associated with both the lateral membranes and the sutures in the lens nucleus (Figure 6B&C, IP3). Repeating these experiments using lenses fixed *in situ* to maintain zonular tension had no effect on the subcellular distribution of AQP0 (data not shown) or AQP5 (Figure 6D) in all regions of interest in the posterior water influx zone. Similarly, pharmacological modulation of zonular tension using tropicamide or pilocarpine had no effect on the distribution of AQP5 in the posterior influx zone (data not shown).

**Figure 6.**
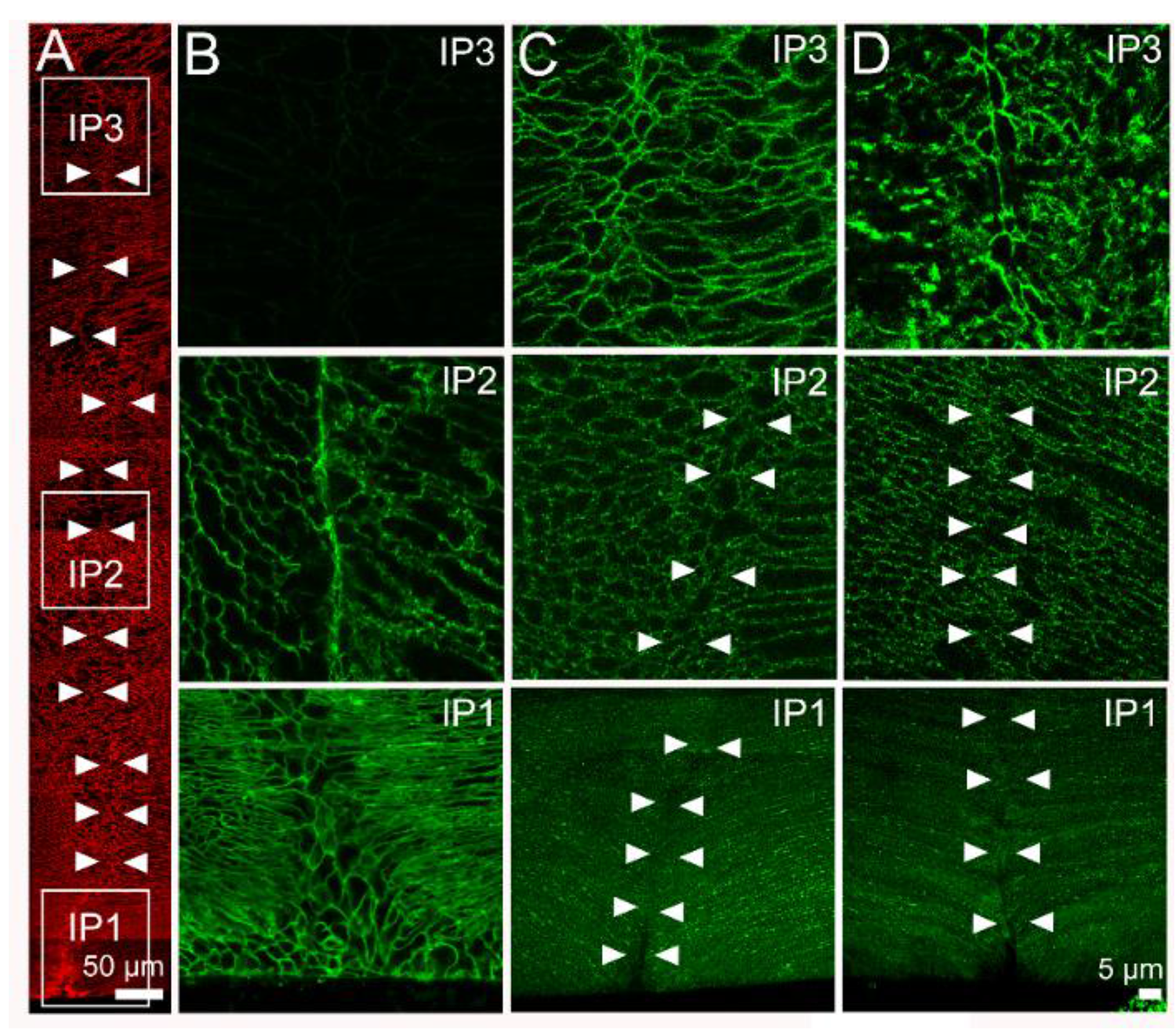
Subcellular localization of lens AQP0 and AQP5 in the influx zone of rat lenses – results from the posterior pole. (A) Image montage of the posterior water influx zone taken from an axial section of a rat lenses that was labelled with the membrane marker WGA (*red*) to highlight suture line (*arrow heads*). Boxes indicate the areas (IP1, IP2 & IP3) from which the higher resolution images shown in images B-D were taken from to investigate the subcellular distribution of AQP0 and AQP5 (*green*). (B) In lenses with cut zonules AQP0 labelling was membranous and strongly labelled the suture in regions IP1 and IP2, but no labelling was observed from region IP3 in the lens core where AQP0 the C-terminus of the AQP0 protein is cleaved. (C) In lenses with zonules cut AQP5 labelling was missing from the suture in regions IP1 and IP2, but labelling was present in the deeper IP3 region. (D) Fixing lenses *in situ* with their zonules attached had no effect on AQP0 (*data not shown*) or AQP5 labelling in the posterior influx zone.

#### AQP distributions along the anterior suture

As shown previously for the efflux zone, AQP1 was only expressed in the epithelial cells that cover the anterior surface of the lens, and hence it was not associated with the anterior suture (Figure 7B, IA1), nor was it expressed in the deeper regions of the lens (Figure 7B, IA2 & IA3). AQP0 labelling around anterior suture was essentially similar to that seen in the posterior suture. In the outer cortex of lenses with zonules cut (Figure 7C) AQP0 was not present in the epithelial cells, was localized to the lateral membranes of fiber cells and was present in the sutures formed from the apical membrane domains (Figure 7C, IA1). AQP0 labelling remained membranous in the fiber cells and strongly localized to the suture in the inner cortex (Figure 7C, IA2). However, as expected AQP0 labelling was not detected in the lens core due to the loss of the antibody epitope (Figure 7C, IA3). AQP5 labelling in the anterior influx zone of lenses with their zonules cut, was essentially similar to that seen in the posterior influx zone. AQP5 labelling was strongly cytoplasmic in the epithelial cells, with a mixed cytoplasmic and membranous localization in the lateral membranes of fiber cells in the outer cortex, and no labelling of the apical membranes associated with the sutures in this region of the lens (Figure 7D, IA1). In the inner cortex AQP5 labelling was associated with the lateral membranes, but was absent from the apical tips at the sutures (Figure 7, IA2), while in the core AQP5 remained membranous in the lateral domains, and was found strongly associated with the sutures (Figure 7D, IA3). However, changes in zonular tension did differentially alter the AQP5 distribution in the anterior influx zone. While no changes in AQP5 labelling were seen in regions IA1 and IA3, of lenses fixed *in situ* with their zonules attached, an increased association of AQP5 with the sutures was observed in region IA2 (Figure 7E). To confirm that a change in zonular tension is the underlying mechanism regulating this observed change in AQP5 labelling in this region of the anterior influx zone, we treated eyes with tropicamide or pilocarpine (Figure 8). We found that in lenses with their zonules attached tropicamide had no effect on AQP5 labelling in the anterior suture (Figure 8B, IA2). However, reducing zonular tension via the application of pilocarpine abolished AQP5 labelling associated with the suture in the inner cortical region of the anterior influx zone (Figure 8C, IA2). In summary, it appears that AQP5 is differentially associated with the apical and basal membrane domains that form anterior and posterior sutures, respectively, in the water influx zone, and in the anterior pole this association with the apical membrane domain can be modified by changes in zonular tension.

**Figure 7.**
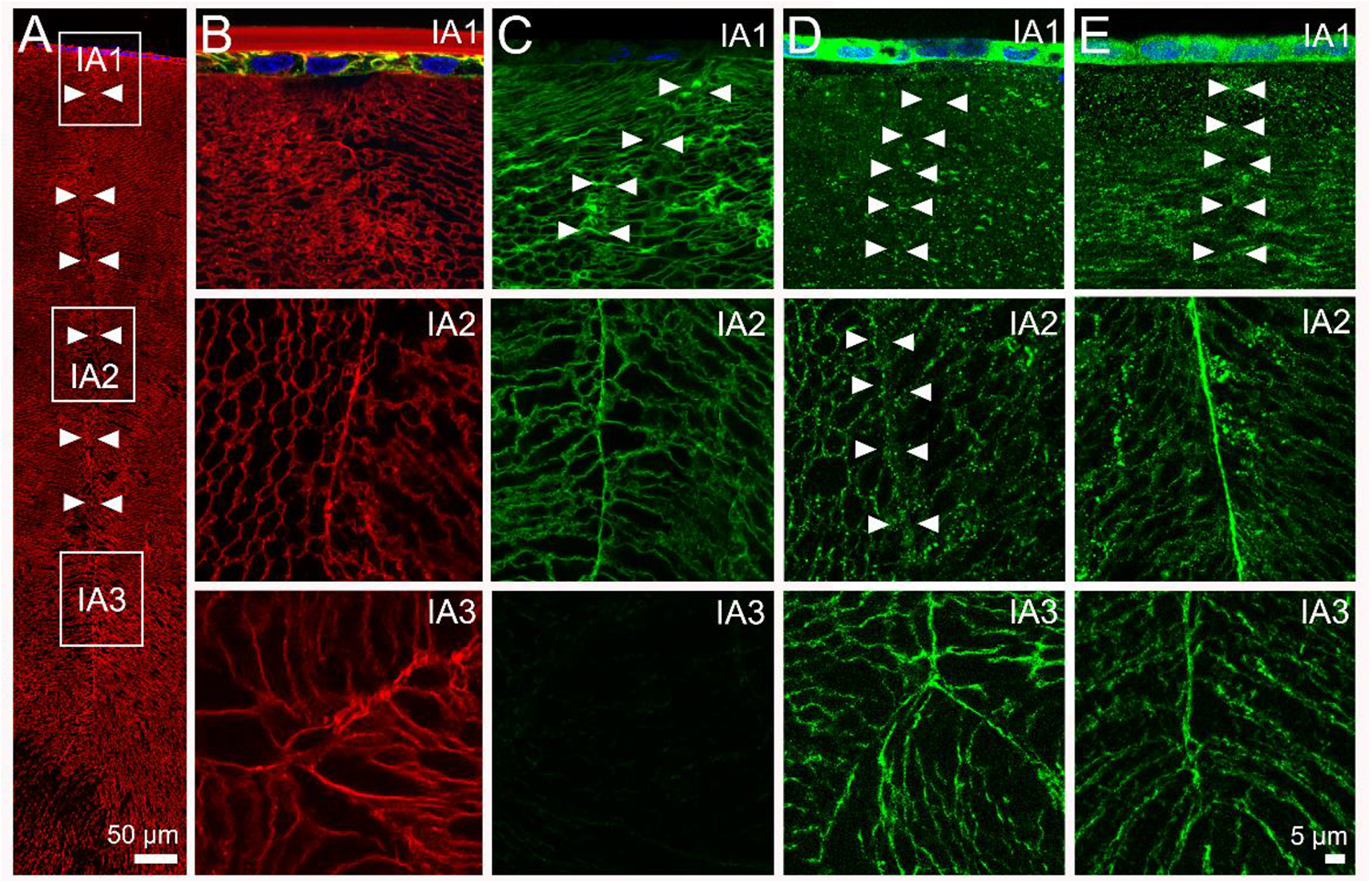
Subcellular localization of lens AQPs in the influx zone of rat lenses – results from the anterior pole. (A) Image montage of the anterior water influx zone taken from an axial section of a rat lenses that was labelled with the membrane marker WGA (*red*) to highlight suture line (*arrow heads*). Boxes indicate the areas (IA1, IA2 & IA3) from which the higher resolution images shown in images B-E were taken from to investigate the subcellular distribution for each lens AQP (*green*). (B) In lenses with cut zonules AQP1 labelling was present only in the epithelial cells of IA1 and was absent from IA2 and IA3. (C) In lenses with cut zonules AQP0 labelling was membranous and strongly labelled the suture in regions IA1 and IA2, but no labelling was observed from region IA3 in the lens core where the C-terminus of AQP0 protein is cleaved. (D) In lenses with zonules cut AQP5 labelling was missing from the suture in regions IA1 and IA2, but labelling was present in the deeper IA3 region. (E) In lenses that were fixed in situ with their zonules attached AQP5 labelling was still absent from the suture in the peripheral region IA1, but was presence in regions IA2 and IA3.

**Figure 8.**
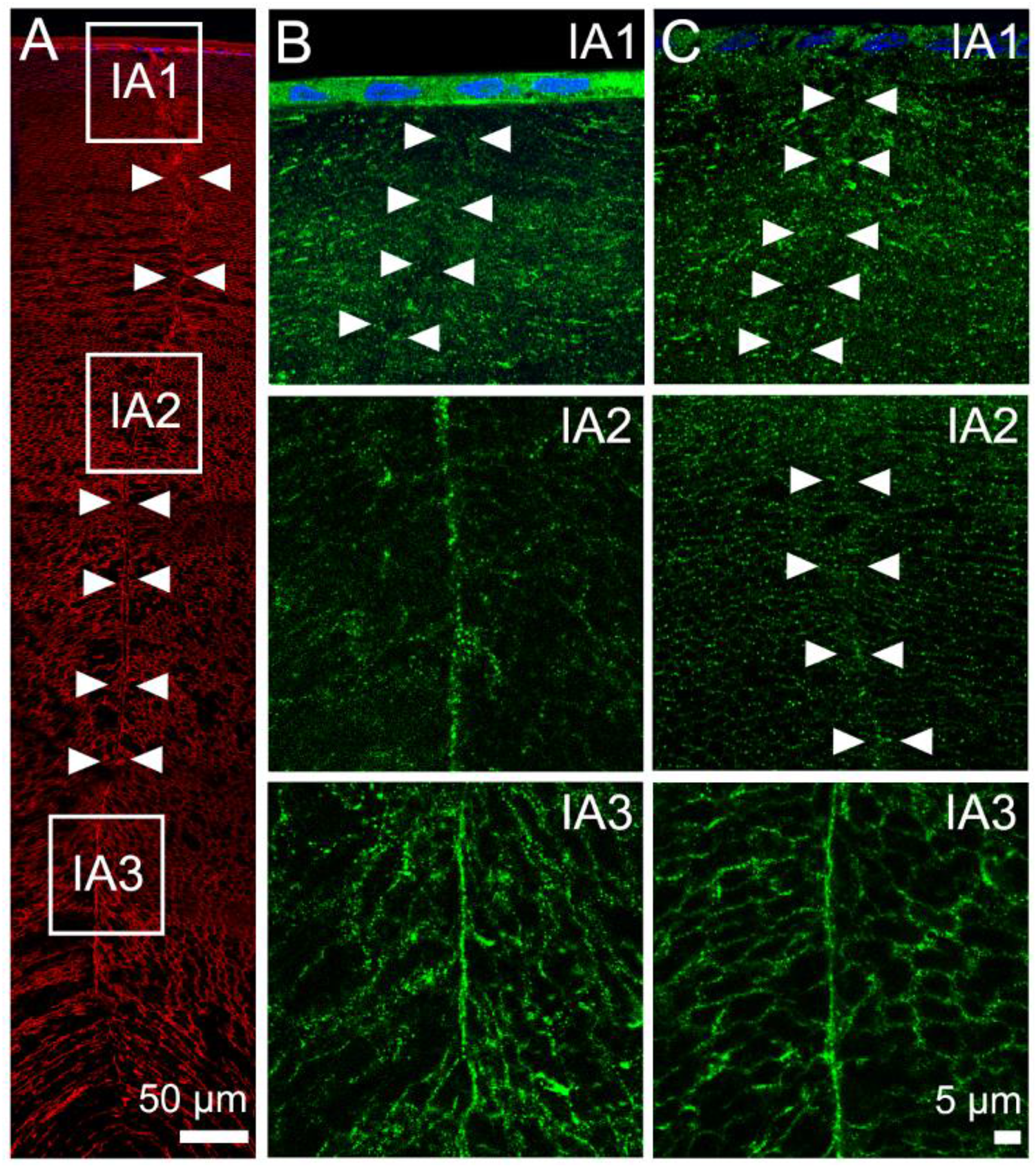
Subcellular distribution of AQP5 in lenses with pharmacologically modulated zonular tension at the anterior suture. (A) Image montage of the anterior water influx zone taken from an axial section of a rat lenses that was labelled with the membrane marker WGA (*red*) to highlight suture line (*arrow heads*). Boxes indicate the areas (IA1, IA2 & IA3) from which the higher resolution images shown in images B&C were taken from to investigate the subcellular distribution of AQP5 (*green*) following the application of tropicamide (B) or pilocarpine for 60 minutes. (B) Application of tropicamide did not change AQP5 labelling in the suture (compare to Figure 6E). (C) In lenses treated with pilocarpine AQP5 labelling was absent from the sutures of regions IA1 and IA2, but was still present at IA3.

## DISCUSSION

Cells move water by actively transporting ions and solutes to establish osmotic gradients that drive the passive diffusion of water through water channels formed from the aquaporin family of proteins^35, 36^. The lens expresses at least three AQPs with very different P_H2O_ and regulation that show spatially distinct patterns of expression and posttranslational modifications. In this study we have utilized immunohistochemistry to visualize the subcellular distribution of the lens AQPs in two spatially discrete areas of the lens that have been shown to be associated water influx and efflux^31, 32^. Our results have confirmed the restriction of AQP1 labelling to the lens epithelium (Figures 2B, 7B), the ubiquitous labelling of AQP0 in the membranes of lens fiber cells (Figures 2C, 6B, 7B), and the loss of AQP0 labelling in the lens nucleus due to the cleavage of the C-terminus of the AQP0 protein (Figure 6B, 7B). In addition, we have shown that these labelling patterns for both AQP1 and AQP0 are unaffected by either mechanical or pharmacological modulation of zonular tension. We have also confirmed the cytoplasmic labelling for AQP5 in epithelial and peripheral fiber cells (Figure 2D) and membrane labelling in deeper fiber cells (Figure 2E), originally observed in equatorial sections obtained from lenses removed from the eye by cutting the zonules^22^. However, in addition we have shown that this labelling pattern is altered by fixing lenses *in situ* to maintain the zonular tension applied to the lens (Figure 3C). In the influx zone we have shown that AQP5 is preferentially localized at the tips of fiber cells that form the anterior (Figure 7D, E), but not the posterior sutures (Figures 6C, D). Furthermore, the membrane localization of AQP5 to the anterior suture was decreased by either the mechanical (Figure 7E, *region A2*) or pharmacological (Figure 8D, *region A2*) reductions in zonular tension applied to the lens. These changes in the subcellular location of AQP5 in response to change in zonular tension, are summarized in Figure 9. Taken together our finding suggest AQP5 acts as a regulated water channel that can dynamically regulate the P_H2O_ of lens fiber cells in the anterior influx and equatorial efflux zones to potentially modulate lens water transport in response to changes in zonular tension.

**Figure 9.**
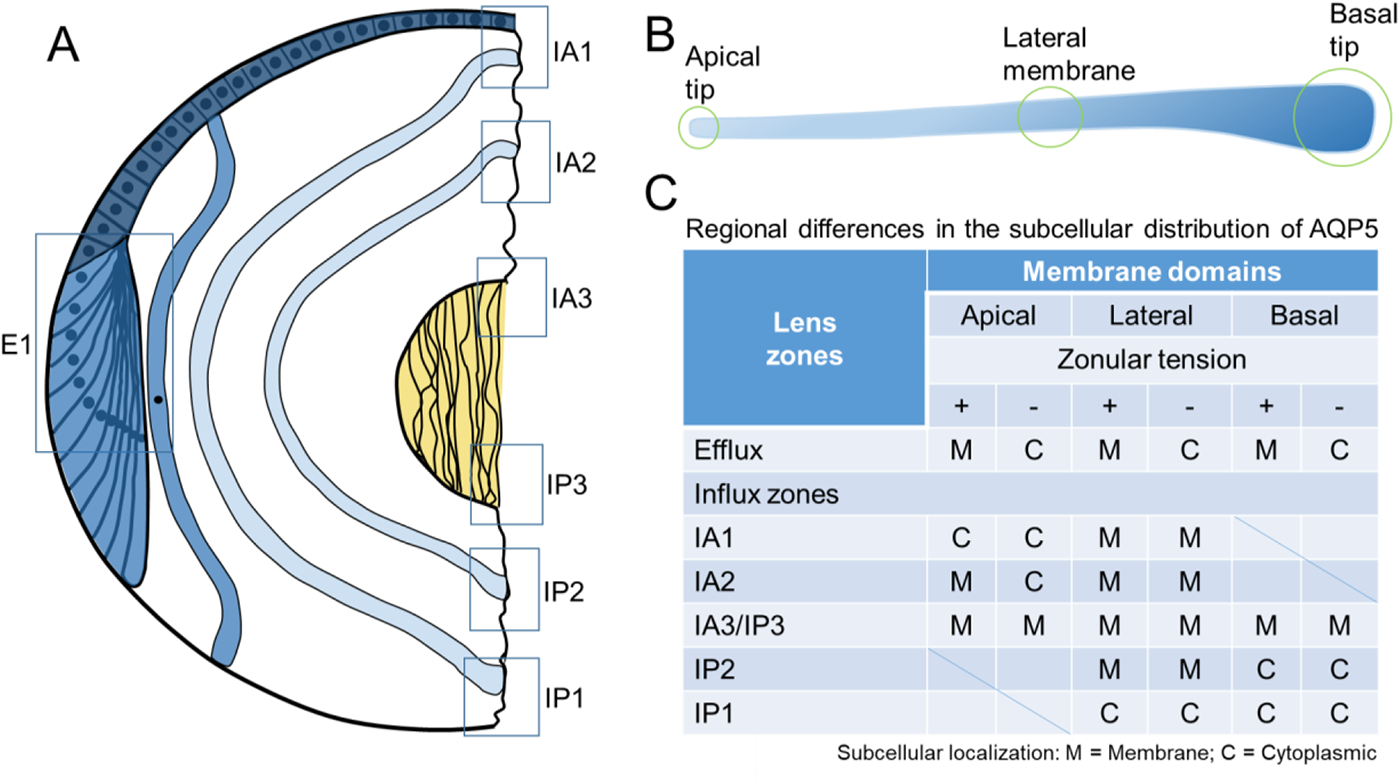
Schematic representation of the subcellular localization of AQP5 in the efflux and influx zones of the rat lens in presence and absence of zonular tension. (A) Epithelial cells (*dark blue*) which differentiate into fiber cells (*blue*) in the equatorial efflux zone (E1) are initially attached by their apical membrane domains to form the modiolus. As fiber cells detach from the modiolus their apical and basal tips migrate along the epithelium and capsule, respectively, and their lateral membranes undergo massive elongation. This process of elongation continues until the apical and basal tips of fiber cells (*light blue*) from opposing lens hemisphere meet to form the anterior and posterior sutures, respectively. As this process continues throughout life newly differentiated secondary fiber cells internalize older mature fiber cells which in turn internalized the primary fiber cells (*yellow*) laid down during embryonic development. Boxes represent the regions in the efflux and influx zones where the subcellular localization of AQP5 was measured. (B) Schematic representation of a fiber cell depicting its three specific membrane domains consisting of the apical tip, lateral membranes and basal tip. (C) Table summarizing the regional differences of AQP5 subcellular localization observed in the efflux and influx zones in the presence and absence of zonular tension.

The existence of a pool of cytoplasmic membrane proteins that translocate to the plasma membrane of fiber cells, either as a function of fiber cell differentiation or dynamically in response to applied stimuli, to confer a specific function required by lens fiber cells, appears to be a reoccurring theme in the lens^37, 38^. In this study, we have shown that in lenses fixed *in situ* the default subcellular location for AQP5 is the membrane, and that upon reducing the zonular tension AQP5 is removed from the membrane, presumably to reduce the P_H2O_ of fiber cell membranes in the anterior and equatorial zones of water influx and efflux, respectively. Interestingly, this removal of AQP5 from the membrane is only transient in lenses organ cultured without their zonules attached (Figure 3B). In an earlier study, we showed a similar increase in AQP5 membrane labelling in the equatorial efflux zone and, in addition, showed that this increase in membrane labelling was associated with an increase in the Hg^+^-sensitive P_H2O_ measured in fiber membrane vesicles derived from organ cultured rat lenses^27^. This earlier study indicates that the changes in the subcellular location of AQP5 labelling observed using immunohistochemistry in the current study are associated with changes in P_H2O_ and support the suggestion that AQP5 functions as a regulated water channel in the anterior influx and equatorial efflux zones of the rat lens.

Despite its potential contributions to the regulation of lens water transport, it was surprising to see that the deletion of the AQP5 gene did not induce a cataract in AQP5-KO lenses^39^. However, it does appear that when AQP5-KO lenses are organ cultured under hyperglycaemic conditions they are more susceptible to the development of cataract, than wild type lenses^39^. This is most probably due to the inability of the AQP5-KO lens to alter its P_H2O_ in response to the osmotic stress induced by the hyperglycaemia, supporting the suggestion that AQP5 plays a role in regulating water fluxes under both steady state conditions and under conditions of stress. Consistent with this idea, an increased expression of AQP5 was observed in epithelial cells removed from patients undergoing cataract surgery^40^. In these patients not only were the expression levels of both AQP5 and AQP1 increased, but AQP5 was found to be more strongly localised to the membranes of lens epithelial cells obtained from cataract patients relative to epithelial cells obtained from age matched non-cataractic lenses. Thus, it appears that as a regulated water channel AQP5 can respond to imposed stresses to modulate water transport in an effort to maintain fluid homeostasis and preserve lens transparency.

In other epithelial tissues, AQP5 has been also shown to act as a regulated water channel that inserts into the apical plasma membrane following phosphorylation of the channel through the cAMP-dependent PKA pathway^23^. In addition, exposure to hypertonic challenge was shown to upregulate AQP5 protein expression through an extracellular signal-regulated kinase (ERK)-dependent pathway in mouse lung epithelial (MLE-15) cells^25^. From the current study, we can add the tension applied to the lens via the zonules as a stimulus capable of regulating the trafficking of AQP5 to the membrane. The observation of this phenomenon raises questions about how changes in zonular tension are sensed and translated to alterations in AQP5 membrane trafficking. Again in other tissues, a synergistic association between the mechano-sensitive TRPV4 and aquaporin water channels has been shown to effect changes in fluid transport to preserve cell volume^41–43^. In the lens, TRPV4 and TRPV1 channels have been shown to not only reciprocally regulate the hydrostatic pressure generated by water flow through gap junction channels^6^, but also to transduce changes to the magnitude of this pressure gradient induced by pharmacologically modulating the zonular tension applied to the lens^7^. Furthermore, changes in zonular tension induced either, mechanically by cutting the zonules^44^, or pharmacologically via application of tropicamide or pilocarpine, also altered the subcellular location of TRPV1 and TRPV4 (data presented at the 6^th^ International Conference on the Lens by Nakazawa *et al*., conference proceedings unpublished). Taken together these observations suggest that in addition to the modulation of the osmotic gradients that drive the transport of water^6, 45^, changes to P_H2O_ are also required to maintain the gradient in hydrostatic pressure that has been measured in all lenses studied to date^5^. Finding a link between TRPV1/4 mediated signalling pathways and the membrane trafficking of AQP5 in lens fiber cells will be a focus of ongoing work.

While similar effects on the membrane localization of AQP5, and presumably the P_H2O_, were evident in the anterior influx and equatorial efflux zones following changes in zonular tension, AQP5 labelling in the posterior influx zone was unaffected, suggesting that changes in zonular tension may differentially affect water influx at the anterior and posterior poles. Structurally the differences in AQP5 localization at the anterior and posterior sutures can be explained by the maintenance of the apical-basal membrane polarity of lens epithelial cells as they differentiate and elongate into fiber cells that then become internalized (Figure 9). The anterior suture is formed by the apical domains of fiber cells that originate from opposing sides of the lens, while the basal domains form the posterior suture^46^. From the current study it would appear that in the outer and inner cortical regions of the lens the apical domains contain AQP5, but the basal domains do not (Figure 6 & 7). Interestingly, in the oldest fiber cells in the lens nucleus this apical-basal polarity is lost and AQP5 is associated with the posterior suture in the nucleus (Figure 7, IP3). This change could be associated with the loss of components of the lens cytoskeleton in the lens nucleus that mediate the anchoring of membrane proteins to specific membrane domains^47–51^.

While it is apparent that the presence of a regulated water channel such as AQP5 can contribute to the maintenance of steady state water transport and water content, the functional significance of the absence of AQP5 in the posterior influx zone is not immediately obvious. However, recent studies performed on zebrafish lenses may provide some insight into the significance of this differential expression AQP5 in the anterior and posterior influx zones. In the zebrafish AQP0 exists in as two functionally divergent isoforms: AQP0a and AQP0b^52–55^. While both AQP0a and AQP0b appear to permeate water, only AQP0b functions as an adhesion molecule^56^. Furthermore, it has been found that the knockout of AQP0a resulted in an anterior polar opacity, suggesting that the water transport function of AQP0a is essential for the formation and maintenance of the anterior suture, but not the posterior suture^55^. Furthermore, in the wild type zebrafish it has been shown that a shift in the location of lens nucleus from an initial anterior position in zebrafish larval to the centre in adult fish is necessary for the development of normal lens optics, and that this process was prevented by the deletion of AQP0a^55^. If we speculate that in the mammalian lens AQP5 and AQP0 are equivalent to zebrafish AQP0a and AQP0b, respectively, then the presence of AQP5 in the anterior suture may be playing a similar role in setting the optical properties of the mammalian lens.

One caveat, however, is that unlike the zebrafish lens, which is round, the mammalian lens has different anterior and posterior radii of curvature. Hence, rather than changing the position (centralisation) of the high refractive index nucleus and therefore lens optical power, as was observed in the developing zebrafish lens, we instead envisage changes to AQP5 mediated water influx at the anterior suture would alter the radius of the curvature of the anterior surface to change the power of the mammalian lens. If this is correct, then the mechanically induced changes in lens shape caused by changes in zonular tension would be amplified by the changes in AQP5 mediated water fluxes in the anterior water influx zone to change the optical power of the lens.

In the rat, which does not appear to accommodate, we envisage that as the lens grows associated changes to zonular tension would produce changes to the steady state optics of the lens to ensure light will remain correctly focussed on the retina. In the human lens, which does accommodate, studies have shown that the anterior curvature of the lens is altered to a greater degree than the posterior curvature^57^. In the accommodated lens, the radius of the anterior lens surface is ~4.7 times greater than the posterior surface, and the anterior surface becomes significantly more hyperbolic^58^. Whether AQP5 plays a role in the process of accommodation in the human lens will be an interesting area to pursue in future work.

In summary, using a series simple immunohistochemical labelling experiments, guided by our knowledge of water transport in the lens, we have shown that a decrease in zonular tension removes AQP5 from the membranes of fibre cells in the equatorial water efflux zone and the anterior water influx zones. Studying how these observations impact the regulation of lens water transport, which is emerging as being essential to the maintenance of the transparent and refractive properties of the lens, has the potential to increase our understand of not only the normal function of the lens, but also how dysfunction of lens water transport can result in lens cataract.

## References

1. Bassnett S, Shi Y, Vrensen GFJM. Biological glass: structural determinants of eye lens transparency. Philosophical Transactions of the Royal Society of London B: Biological Sciences 2011;366:1250–1264.

2. Kuszak JR, Zoltoski RK, Tiedemann CE. Development of lens sutures. Int J Dev Biol 2004;48:889–902.

3. Mathias RT, Kistler J, Donaldson P. The lens circulation. Journal of Membrane Biology 2007;216:1–16.

4. Gao J, Sun X, Moore LC, White TW, Brink PR, Mathias RT. Lens intracellular hydrostatic pressure is generated by the circulation of sodium and modulated by gap junction coupling. The Journal of General Physiology 2011;137:507–520.

5. Gao J, Sun X, Moore LC, Brink PR, White TW, Mathias RT. The effect of size and species on lens intracellular hydrostatic pressure. Investigative Ophthalmology & Visual Science 2013;54:183.

6. Gao J, Sun X, White TW, Delamere NA, Mathias RT. Feedback Regulation of Intracellular Hydrostatic Pressure in Surface Cells of the Lens. Biophysical Journal 2015;109:1830–1839.

7. Chen Y, Gao J, Li L, et al. The Ciliary Muscle and Zonules of Zinn Modulate Lens Intracellular Hydrostatic Pressure Through Transient Receptor Potential Vanilloid Channels. Investigative ophthalmology & visual science 2019;60:4416–4424.

8. Kozono D, Yasui M, King LS, Agre P. Aquaporin water channels: atomic structure molecular dynamics meet clinical medicine. The Journal of Clinical Investigation 2002;109:1395–1399.

9. Varadaraj K, Kushmerick C, Baldo GJ, Bassnett S, Shiels A, Mathias RT. The role of MIP in lens fiber cell membrane transport. The Journal of Membrane Biology 1999;170:191–203.

10. Grey AC, Walker KL, Petrova RS, et al. Verification and spatial localization of aquaporin-5 in the ocular lens. Experimental Eye Research 2013;108:94–102.

11. Ruiz-Ederra J, Verkman AS. Accelerated cataract formation and reduced lens epithelial water permeability in aquaporin-1-deficient mice. Investigative Ophthalmology & Visual Science 2006;47:3960–3967.

12. Varadaraj K, Kumari SS, Mathias RT. Functional expression of aquaporins in embryonic, postnatal, and adult mouse lenses. Developmental Dynamics 2007;236:1319–1328.

13. Bok D, Dockstader J, Horwitz J. Immunocytochemical localization of the lens main intrinsic polypeptide (MIP26) in communicating junctions. The Journal of Cell Biology 1982;92:213–220.

14. Yang B, Verkman AS. Water and glycerol permeabilities of aquaporins 1-5 and MIP determined quantitatively by expression of epitope-tagged constructs in Xenopus oocytes. Journal of Biological Chemistry 1997;272:16140–16146.

15. Ball LE, Little M, Nowak MW, Garland DL, Crouch RK, Schey KL. Water permeability of C-terminally truncated aquaporin 0 (AQP0 1-243) observed in the aging human lens. Investigative Ophthalmology & Visual Science 2003;44:4820–4828.

16. Lo W-K, Harding CV. Square arrays and their role in ridge formation in human lens fibers. Journal of ultrastructure research 1984;86:228–245.

17. Zampighi GA, Hall JE, Ehring GR, Simon SA. The structural organization and protein composition of lens fiber junctions. The Journal of Cell Biology 1989;108:2255–2275.

18. Gonen T, Cheng Y, Sliz P, et al. Lipid-protein interactions in double-layered two-dimensional AQP0 crystals. Nature 2005;438:633–638.

19. Rose KML, Gourdie RG, Prescott AR, Quinlan RA, Crouch RK, Schey KL. The C terminus of lens aquaporin 0 interacts with the cytoskeletal proteins filensin and CP49. Investigative Ophthalmology & Visual Science 2006;47:1562–1570.

20. Wang Z, Schey KL. Aquaporin-0 interacts with the FERM domain of ezrin/radixin/moesin proteins in the ocular lens. Investigative Ophthalmology & Visual Science 2011;52:5079.

21. Grey AC, Li L, Jacobs MD, Schey KL, Donaldson PJ. Differentiation-dependent modification and subcellular distribution of aquaporin-0 suggests multiple functional roles in the rat lens. Differentiation 2009;77:70–83.

22. Petrova RS, Schey KL, Donaldson PJ, Grey AC. Spatial distributions of AQP5 and AQP0 in embryonic and postnatal mouse lens development. Experimental Eye Research 2015;132:124–135.

23. Woo J, Chae YK, Jang SJ, et al. Membrane trafficking of AQP5 and cAMP dependent phosphorylation in bronchial epithelium. Biochemical and biophysical research communications 2008;366:321–327.

24. Ishikawa Y, Eguchi T, Skowronski MT, Ishida H. Acetylcholine acts on M 3 muscarinic receptors and induces the translocation of aquaporin5 water channel via cytosolic Ca 2+ elevation in rat parotid glands. Biochemical and Biophysical Research Communications 1998;245:835–840.

25. Hoffert JD, Leitch V, Agre P, King LS. Hypertonic induction of aquaporin-5 expression through an ERK-dependent pathway. Journal of Biological Chemistry 2000;275:9070–9077.

26. Moore M, Ma T, Yang B, Verkman A. Tear secretion by lacrimal glands in transgenic mice lacking water channels AQP1, AQP3, AQP4 and AQP5. Experimental eye research 2000;70:557–562.

27. Petrova RS, Webb KF, Vaghefi E, Walker K, Schey KL, Donaldson PJ. Dynamic functional contribution of the water channel AQP5 to the water permeability of peripheral lens fiber cells. American Journal of Physiology-Cell Physiology 2017;314:ajpcell.00214.02017.

28. Wang W-H, Millar JC, Pang I-H, Wax MB, Clark AF. Noninvasive measurement of rodent intraocular pressure with a rebound tonometer. Investigative ophthalmology & visual science 2005;46:4617–4621.

29. Bassnett S. A method for preserving and visualizing the three-dimensional structure of the mouse zonule. Experimental eye research 2019;185:107685.

30. Jacobs MD, Donaldson PJ, Cannell MB, Soeller C. Resolving morphology and antibody labeling over large distances in tissue sections. Microscopy Research and Technique 2003;62:83–91.

31. Candia OA, Mathias R, Gerometta R. Fluid circulation determined in the isolated bovine lens. Investigative Ophthalmology & Visual Science 2012;53:7087.

32. Vaghefi E, Pontre BP, Jacobs MD, Donaldson PJ. Visualizing ocular lens fluid dynamics using MRI: manipulation of steady state water content and water fluxes. American Journal of Physiology-Regulatory, Integrative and Comparative Physiology 2011;301:R335–R342.

33. Zampighi GA, Simon SA, Hall JE. The specialized junctions of the lens. Internaional Review of Cytology 1992;136:185–225.

34. Al-Ghoul KJ, Lindquist TP, Kirk SS, Donohue ST. A Novel Terminal Web-Like Structure in Cortical Lens Fibers: Architecture and Functional Assessment. The Anatomical Record: Advances in Integrative Anatomy and Evolutionary Biology 2010;293:1805–1815.

35. Verkman AS, Ruiz-Ederra J, Levin MH. Functions of aquaporins in the eye. Progress in Retinal and Eye Research 2008;27:420–433.

36. Schey KL, Wang Z, Wenke JL, Qi Y. Aquaporins in the eye: expression, function, and roles in ocular disease. Biochimica et Biophysica Acta (BBA)-General Subjects 2014;1840:1513–1523.

37. Donaldson PJ, Grey AC, Merriman-Smith BR, et al. Functional imaging: new views on lens structure and function. Clinical and Experimental Pharmacology and Physiology 2004;31:890–895.

38. Donaldson PJ, Lim J. Membrane transporters. Ocular Transporters in Ophthalmic Diseases and Drug Delivery: Springer; 2008:89–110.

39. Kumari SS, Varadaraj K. Aquaporin 5 knockout mouse lens develops hyperglycemic cataract. Biochemical and biophysical research communications 2013;441:333–338.

40. Barandika O, Ezquerra-Inchausti M, Anasagasti A, et al. Increased aquaporin 1 and 5 membrane expression in the lens epithelium of cataract patients. Biochimica et Biophysica Acta (BBA)-Molecular Basis of Disease 2016;1862:2015–2021.

41. Liu X, Bandyopadhyay B, Nakamoto T, et al. A Role for AQP5 in Activation of TRPV4 by Hypotonicity CONCERTED INVOLVEMENT OF AQP5 AND TRPV4 IN REGULATION OF CELL VOLUME RECOVERY. Journal of Biological Chemistry 2006;281:15485–15495.

42. Benfenati V, Caprini M, Dovizio M, et al. An aquaporin-4/transient receptor potential vanilloid 4 (AQP4/TRPV4) complex is essential for cell-volume control in astrocytes. Proceedings of the National Academy of Sciences 2011;108:2563–2568.

43. Taguchi D, Takeda T, Kakigi A, Takumida M, Nishioka R, Kitano H. Expressions of aquaporin-2, vasopressin type 2 receptor, transient receptor potential channel vanilloid (TRPV) 1, and TRPV4 in the human endolymphatic sac. The Laryngoscope 2007;117:695–698.

44. Nakazawa Y, Donaldson PJ, Petrova RS. Verification and spatial mapping of TRPV1 and TRPV4 expression in the embryonic and adult mouse lens. Experimental eye research 2019;186:107707.

45. Shahidullah M, Mandal A, Delamere NA. Activation of TRPV1 channels leads to stimulation of NKCC1 cotransport in the lens. American Journal of Physiology-Cell Physiology 2018;315:C793–C802.

46. Zampighi GA, Eskandari S, Kreman M. Epithelial organization of the mammalian lens. Experimental Eye Research 2000;71:415–435.

47. Sandilands A, Prescott AR, Carter J, et al. Vimentin and CP49/filensin form distinct networks in the lens which are independently modulated during lens fibre cell differentiation. Journal of cell science 1995;108:1397–1406.

48. Blankenship TN, Hess JF, FitzGerald PG. Development-and differentiation-dependent reorganization of intermediate filaments in fiber cells. Investigative ophthalmology & visual science 2001;42:735–742.

49. FitzGerald PG. Lens intermediate filaments. Experimental eye research 2009;88:165–172.

50. Oka M, Kudo H, Sugama N, Asami Y, Takehana M. The function of filensin and phakinin in lens transparency. Molecular vision 2008;14:815.

51. Xu L, Overbeek PA, Reneker LW. Systematic analysis of E-, N-and P-cadherin expression in mouse eye development. Experimental eye research 2002;74:753–760.

52. Froger A, Németh-Cahalan K, Kalman K, Schilling TF, Hall JE. Knockdown of Zeb1-AQP0 or Zeb2-AQP0 Leads to Cataract Formation in Zebrafish. Investigative Ophthalmology & Visual Science 2008;49:3170–3170.

53. Froger A, Clemens D, Kalman K, Németh-Cahalan KL, Schilling TF, Hall JE. Two distinct aquaporin 0s required for development and transparency of the zebrafish lens. Investigative Ophthalmology & Visual Science 2010;51:6582.

54. Clemens DM, Németh-Cahalan KL, Trinh LT, Zhang T, Schilling TF, Hall JE. In Vivo Analysis of Aquaporin 0 Function in Zebrafish: Permeability Regulation Is Required for Lens TransparencyIn Vivo Analysis of Aquaporin. Investigative Ophthalmology & Visual Science 2013;54:5136–5143.

55. Vorontsova I, Gehring I, Hall JE, Schilling TF. Aqp0a regulates suture stability in the zebrafish lens. Investigative ophthalmology & visual science 2018;59:2869–2879.

56. Vorontsova I, Vallmitjana A, Nakazawa Y, et al. Aquaporin 0A is Required for Water Homeostasis in the Zebrafish Lens In Vivo. Biophysical journal 2020;118:167a.

57. Dubbelman M, Van der Heijde GL. The shape of the aging human lens: curvature, equivalent refractive index and the lens paradox. Vision research 2001;41:1867–1877.

58. Dubbelman M, Van der Heijde GL, Weeber HA. Change in shape of the aging human crystalline lens with accommodation. Vision research 2005;45:117–132.

